# Positive memory specificity reduces adolescent vulnerability to depression

**DOI:** 10.1101/329409

**Authors:** Adrian Dahl Askelund, Susanne Schweizer, Ian M. Goodyer, Anne-Laura van Harmelen

## Abstract

Depression is the leading cause of ill health and disability worldwide^1^. A known risk factor of depression is exposure to early life stress^2^. Such early stress exposure has been proposed to sensitise the maturing psychophysiological stress system to later life stress^3^. Activating positive memories dampens acute stress responses with resultant lower cortisol response and improved mood in humans^4^ and reduced depression-like behaviour in mice^5^. It is unknown whether recalling positive memories similarly reduces adolescent vulnerability to depression. Here we used path modelling to examine the effects of positive autobiographical memory specificity on later morning cortisol and negative self-cognitions during low mood in adolescents at risk for depression due to early life stress (n = 427, age: 14 years)^6^. We found that experimentally assessed positive but not negative memory specificity was associated with lower morning cortisol and less negative self-cognitions during low mood one year later. Moderated mediation analyses demonstrated that positive memory specificity reduced later depressive symptoms through lowering negative self-cognitions in response to negative life events reported in the one-year interval. Positive memory specificity actively dampened the negative effect of stressors over time, thereby operating as a resilience factor reducing the risk of subsequent depression^7^. These findings suggest that developing methods to improve positive memory specificity in at-risk adolescents may counteract vulnerability to depression.

---

Remembering specific positive life experiences, as single, temporally limited instances from the past, may be an important protective process when stress occurs^4^. People engage in reminiscing about past events quite frequently in their everyday lives^8^, and evidence suggests that healthy individuals use recall of positive memories as one of many strategies to repair sad mood^9^. Positive emotions, for instance generated by such memories, in turn appear to facilitate physiological and emotional stress recovery, particularly in resilient individuals^10,11^. Recalling positive memories may be a protective mechanism in most adolescents, which may be disturbed in individuals who are vulnerable to depression^12^. In support of this, adolescents who were in remission from a recent depressive episode recalled more categorical memories^13^. Furthermore, it was recently found that depressed, at-risk and healthy adolescents show a gradient of positive memory deficits after a negative mood induction^14^. These findings together imply that less specific responses to positive cues in particular (‘positive memory specificity’) constitute a trait-like marker of depressive vulnerability in at-risk adolescents. In addition, having a tendency toward more categorical, overgeneral (i.e., lacking in defining characteristics) memories that are not fixed in time or place, is characteristic of depression^15^. Low memory specificity is a trait-like characteristic of individuals at risk for depression^16,17^, those currently depressed^13^, and those in remission from depression^18^. Crucially, low memory specificity predicts the onset and course of depression^18^, especially in response to stress^19^. Thus, low memory specificity may comprise a cognitive mechanism through which stress increases the risk of developing depression. Here we examined whether positive memory specificity is related to lower cognitive and physiological vulnerability to depression cross-sectionally and prospectively in adolescents at risk due to high emotionality and/or exposure to early life stress.

We examined the effect of positive memory specificity on two types of vulnerability for depression: negative self-cognitions during low mood^20^–^23^ and high morning cortisol^6,17,24,25^. Negative self-cognitions refer to the tendency to blame and be derogatory about oneself (“I am useless”). Negative self-cognitions can be reactivated during in stress in individuals who are in remission from depression^23^ and have been shown to prospectively predict first incidence of depression^26^. In individuals at risk for depression with a negative thinking style, negative life events may be particularly detrimental. The capacity to recall positive memories, however, may attenuate the interactive risks conferred by stress-exposure and negative self-cognitions. Morning cortisol is a physiological marker of vulnerability to depression; high morning cortisol is associated with familial risk for^25^, onset^6,17^, presence^24^ and history of^24^ major depression. Recently, morning cortisol was shown to interact with stressful life events leading to more depressive symptoms in adolescent girls^27^. Recalling positive memories, in contrast, has been shown to dampen the cortisol response to stress^4^. Here, we therefore hypothesised that positive memory specificity would be associated with less negative self-cognitions during low mood and lower morning cortisol over time. That is, we investigated the putative effects of positive memory specificity on two distinct vulnerability pathways for depression; one cognitive and the other physiological^28^.

In this study, the role of positive memory specificity was investigated prospectively in a sample of adolescents at-risk for depression due to early life stress and/or high emotionality. Here, early life stress was operationalised as the presence of any early risk factor including current marital disharmony or past breakdown, moderately to severely negative life events, parental psychiatric illness, and/or the loss of a close relative or friend. In this letter, we use the term more broadly when referring to studies that examined childhood emotional, physical or sexual abuse and/or neglect. High emotionality was defined as scoring over the 80^th^ percentile on this trait^29^. All participants (n = 427, 200 girls, age 14; see descriptive statistics in Supplementary Table 1) completed the experimental cued recall Autobiographical Memory Test at baseline^30^. We used the ratio of total specific divided by total categorical (overgeneral) responses to positive cues as our predictor variable. The rationale for using this ratio was that specific and categorical responses are thought to tap into the same underlying construct of positive memory specificity (see Supplementary Results for a validation of this ratio). At baseline and 1-year follow-up, all participants reported the frequency of moderate to severe negative life events during the last 12 months in a semi-structured interview. At both times, all participants reported depressive symptoms during the last two weeks (Mood and Feelings Questionnaire^31^), and negative self-cognitions and dysphoric mood experiences during episodes of low mood in the past month^23^. In accordance with Teasdale’s Differential Activation hypothesis^23^, we used the ratio of negative self-cognitions divided by dysphoric mood as our measure of cognitive vulnerability to depression. To acquire a stable trait-like measure of morning cortisol, a latent factor was extracted from morning cortisol across four sampling days at both baseline and follow-up (see Supplementary Results and Supplementary Figure 1). The morning cortisol factor showed strong measurement invariance over time, therefore, changes in cortisol can be meaningfully interpreted (see Supplementary Table 2).

We used path modelling in R (*lavaan*^32^) with a robust estimator to examine whether positive memory specificity was related to less negative self-cognitions during low mood and lower morning cortisol currently and/or one year later. IQ and gender were specified as covariates since they have been associated with cognitive and physiological vulnerability for depression^17,33^. We also included negative life events as a covariate in the model because we were interested in depressive vulnerability relative to the extent of exposure to recent life stress^34^. We found that positive memory specificity at baseline was related to less negative self-cognitions during low mood at follow-up (Pearson’s r effect size = -.14, 95 % CI [-.23, - .05], P = .003), but not at baseline (r = -.05, 95 % CI [-.14, .05], P = .299). Positive memory specificity was also related to lower morning cortisol at follow-up (r = -.13, 95 % CI [-.22, - .04], P = .006), but not at baseline (r = -.09, 95 % CI [-.18, .004], P = .064). Model fit was excellent (see Figure 1 and Table 1). The findings were not influenced by outliers (see Supplementary Table 3) or selective attrition (see Supplementary Table 4). The absence of cross-sectional relations was not due to the inclusion of follow-up assessments in the model, as post hoc analyses showed no significant raw correlations between positive memory specificity and baseline cortisol (r = -.08, 95 % CI [-.17, .02], P = .103) or negative self-cognitions during low mood (r = -.06, 95 % CI [-.15, .04], P = .252).

**Figure 1.**
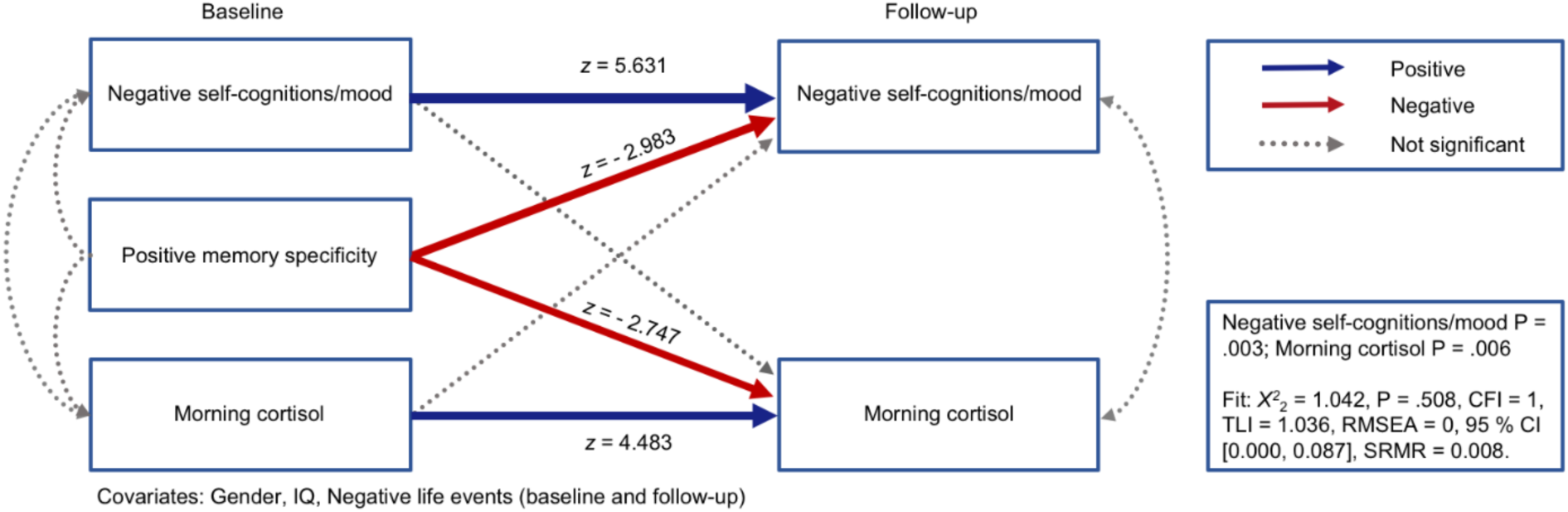
Positive memory specificity predicts lower cognitive and physiological vulnerability at follow-up. Path model showing that positive memory specificity is related to both lower negative self-cognitions during low mood and morning cortisol at follow-up, and only morning cortisol at baseline. Broader arrows indicate stronger relationships. *z*-value = standardised path coefficient.

**Table 1.** Positive memory specificity predicts lower negative self-cognitions during low mood and morning cortisol. n = 427. (b) = baseline, (f) = follow-up. Boys are coded as 1, girls as 2. Significant paths are bolded. Robust model fit indices: *X*^2^_2_ = 1.042, P = .508, CFI = 1, TLI = 1.036, RMSEA = 0, 90 % CI [0.000, 0.087], SRMR = 0.008. Estimate = unstandardised path coefficient, S.E. = robust standard error, *z*-value = standardised path coefficient.

| Outcome | Predictor | Estimate | S.E. | z-value | P(>\|z\|) |
| --- | --- | --- | --- | --- | --- |
| **Morning cortisol (b)** | Positive memory specificity (b) | -0.305 | 0.165 | -1.851 | .064 |
|  | Negative life events (b) | 0.012 | 0.060 | 0.198 | .843 |
|  | Gender (b) | 0.677 | 0.115 | 5.878 | **.001** |
|  | IQ (b) | -0.000 | 0.003 | -0.087 | .931 |
| **Morning cortisol (f)** | Morning cortisol (b) | 0.363 | 0.081 | 4.483 | **.001** |
|  | Positive memory specificity (b) | -0.360 | 0.131 | -2.747 | **.006** |
|  | Negative self-cognitions/mood (b) | 0.144 | 0.137 | 1.054 | .292 |
|  | Negative life events (b) | 0.008 | 0.053 | 0.156 | .876 |
|  | Negative life events (f) | 0.083 | 0.048 | 1.726 | .084 |
|  | Gender (b) | 0.288 | 0.106 | 2.730 | **.006** |
|  | IQ (b) | 0.011 | 0.003 | 3.772 | **.001** |
| **Negative self-cognitions/mood (b)** | Positive memory specificity (b) | -0.048 | 0.046 | -1.038 | .299 |
|  | Negative life events (b) | 0.022 | 0.016 | 1.433 | .152 |
|  | Gender (b) | 0.032 | 0.032 | 1.002 | .317 |
|  | IQ (b) | -0.001 | 0.001 | -0.802 | .423 |
| **Negative self-cognitions/mood (f)** | Negative self-cognitions/mood (b) | 0.399 | 0.071 | 5.631 | **.001** |
|  | Positive memory specificity (b) | -0.115 | 0.039 | -2.983 | **.003** |
|  | Morning cortisol (b) | -0.012 | 0.012 | -0.978 | .328 |
|  | Negative life events (b) | 0.015 | 0.012 | 1.288 | .198 |
|  | Negative life events (f) | 0.015 | 0.013 | 1.180 | .238 |
|  | Gender (b) | 0.019 | 0.030 | 0.627 | .531 |
|  | IQ (b) | 0.000 | 0.001 | 0.512 | .609 |
| **Morning cortisol (b) ~** | Negative self-cognitions/mood (b) | 0.026 | 0.019 | 1.370 | .171 |
| **Morning cortisol (f) ~** | Negative self-cognitions/mood (f) | 0.000 | 0.013 | 0.036 | .972 |

Next, we examined whether the effects in the path model (Figure 1 and Table 1) were due to memory specificity in general (and also found for negative memory specificity), or specific to positive memory specificity. We ran an exploratory model with both negative and positive memory specificity as predictors. In this model, there was a relation between positive memory specificity and negative self-cognitions/mood (Effect = -0.122, S.E. = 0.041, *z* = -2.979, Pearson’s r effect size = -.14, 95 % CI [-.23, -.05], P = .003) and morning cortisol at follow-up (Effect = -0.368, S.E. = 0.146, *z* = -2.523, r = -.12, 95 % CI [-.21, -.03], P = .012). In contrast, negative memory specificity was unrelated to negative self-cognitions/mood (Effect = 0.018, S.E. = 0.043, *z* = 0.422, r = .02, 95 % CI [-.08, .11], P = .673) and morning cortisol at follow-up (Effect = 0.021, S.E. = 0.153, *z* = 0.134, r = .01, 95 % CI [-.08, .10], P = .893). Relationships between positive memory specificity and negative self-cognitions/mood (Effect = -0.033, S.E. = 0.049, *z* = -0.649, r = -.03, 95 % CI [-.12, .07], P = .497) and morning cortisol were not significant at baseline (Effect = -0.263, S.E. = 0.179, *z* = -1.469, r = -.07, 95 % CI [- .16, .03], P = .142). Negative memory specificity was unrelated to negative self-cognitions/mood (Effect = -0.038, S.E. = 0.049, *z* = -0.774, r = -.04, 95 % CI [-.13, .06], P = .439) and morning cortisol at baseline (Effect = -0.108, S.E. = 0.169, *z* = -0.640, r = -.03, 95 % CI [-.12, .07], P = .522). Robust fit statistics indicated good fit for the model with both predictors (*X*^2^_2_ = 1.361, P = .506, CFI = 1, TLI = 1.041, RMSEA = 0, 95 % CI [0.000, 0.087], SRMR = 0.007). In this model, constraining the negative memory specificity paths to zero did not affect model fit, suggesting that negative memory specificity was not needed to explain our data (ANOVA: *X*^*2*^_*4*_ = 1.601, P = .910; robust fit statistics still indicated good fit: *X*^2^_4_ = 1.558, P = .816, CFI = 1, TLI = 1.078, RMSEA = 0, 95 % CI [0.000, 0.045], SRMR = 0.008). On the other hand, constraining the positive memory specificity paths to zero significantly lowered model fit (ANOVA: *X*^*2*^_*4*_ = 16.659, P < .001; robust fit statistics indicated poor model fit: *X*^2^_4_ = 16.869, P = .002, CFI = 0.947, TLI = 0.605, RMSEA = .086, 95 % CI [0.047, 0.131], SRMR = 0.020). Furthermore, the lack of an effect of negative memory specificity was not due to the inclusion of positive memory specificity in the same model. When positive memory specificity was constrained to zero, negative memory specificity was unrelated to negative self-cognitions/mood (Effect = -0.035, S.E. = 0.041, *z* = -0.844, r = -.04, 95 % CI [- .13, .06], P = .399) and morning cortisol at follow-up (Effect = -0.139, S.E. = 0.136, *z* = -1.020, r = -.05, 95 % CI [-.14, .05], P = .308). Overall, positive but not negative memory specificity contributed to the path model, so negative memory specificity was not needed as a predictor.

Accessing specific positive memories in the face of stress may activate a cognitive mechanism that ‘disconfirms’ negative self-cognitions, leading indirectly to mood improvement over time. To test this mechanistic hypothesis, we first ran a moderation analysis with prospective negative life events as a moderator of the relationship between positive memory specificity at baseline and negative self-cognitions at follow-up. We conducted a bias-corrected moderation analysis with 5,000 bootstrapped samples using the PROCESS macro in SPSS^35^. This analysis supported our hypothesis (see Table 2 and Supplementary Figure 2), showing a significant overall moderation (F_1,419_ = 7.927, P = .005), controlling for negative self-cognitions at baseline. Positive memory specificity predicted lower negative self-cognitions in those who experienced at least one negative life event (Pearson’s r effect size = -.21, 95 % CI [-.30, -.12]), but not in those who did not experience any negative life events (r = -.05, 95 % CI [-.14, .05]). In contrast, post hoc analyses showed that negative life events did not moderate the relationship between positive memory specificity and dysphoric mood (F_1,419_ = 1.785, P = .182) depressive symptoms (F_1,419_ = 1.534, P = .216) or morning cortisol (F_1,419_ = 0.271, P = .603) at follow-up, controlling for baseline values. Next, we explored whether negative self-cognitions mediated an indirect relationship between positive memory specificity and later depressive symptoms depending on exposure to negative life events (i.e., a moderated mediation; Figure 2). In line with the path model in Figure 1, we controlled for baseline depressive symptoms and negative self-cognitions in the moderated mediation analysis to focus on differences over time. This analysis (see Table 2 and Figure 2) showed a significant indirect effect of positive memory specificity through lower negative self-cognitions on depressive symptoms, depending on exposure to negative life events (Index = -3.026, S.E. = 1.290, 95 % CI [-5.752, -0.704]).

**Figure 2.**
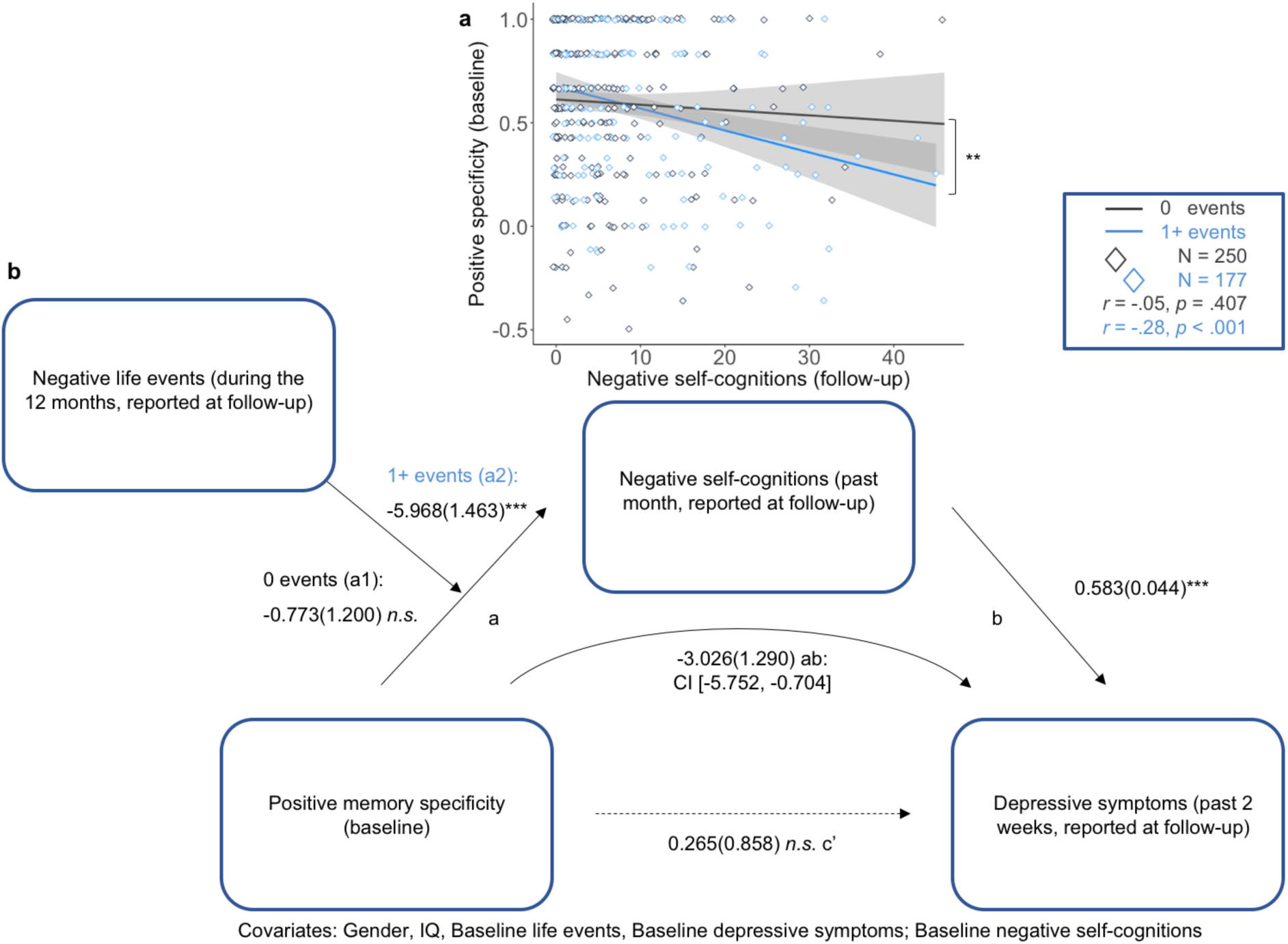
Positive memory specificity lowers depressive symptoms after recent negative life events.

n = 427. Plot **a** is showing a significant interaction where the effect of positive memory specificity on negative self-cognitions depends on exposure to recent negative life events. Specifically, positive memory specificity is moderately related to lower negative self-cognitions in those exposed to one or more recent negative life events (during the 12 months of the study; blue line). The relationship is small and not significant in those not exposed to recent negative life events (black line). Lines show raw correlations, and grey bands show confidence intervals. Figure **b** shows a moderated mediation model where positive memory specificity at baseline decreases depressive symptoms indirectly over time. Its effect is mediated by less negative self-cognitions, depending upon exposure to negative life events. *Path a:* Effect of positive memory specificity on negative self-cognitions, depending on exposure to recent negative life events; *path b:* relationship between negative self-cognitions and depressive symptoms; *path c’:* Effect of positive memory specificity at baseline on depressive symptoms at follow-up, controlling for the indirect effect; *path ab:* the index of the conditional indirect effect of positive memory specificity on depressive symptoms. The 95 % confidence interval (CI) for this indirect path does not include 0, suggesting that the moderated mediation is significantly different from 0 (at P < ,05). Path values represent unstandardised coefficients and bootstrapped standard errors; *P < .05; **P < .01; ***P < .005; *n.s.* not significant.

The moderation model showed the same results without any covariates (F_1,423_ = 8.039, P = .005; see Supplementary Table 5) and with outliers excluded (F_1,382_ = 6.755, P = .010; see Supplementary Table 6). Also, the moderated mediation model showed the same results without any covariates (Index = -4.788, S.E. = 1.859, 95 % CI [-8.541, -1.255]; see Supplementary Table 5) and with outliers excluded (Index = -2.206, S.E. = 1.034, 95 % CI [- 4.301, -0.291]; see Supplementary Table 6). Importantly, the moderated mediation model was specified on data from two and not three waves (see correlations between the cross-sectional measures in the model in Supplementary Results). However, a moderated mediation model with the mediator and outcome interchanged showed that depressive symptoms did not mediate the effect of positive memory specificity on negative self-cognitions (Index = -1.184, S.E. = 1.167, 95 % CI [-3.630, 0.962]; see Table 2).

**Table 2.** Results of moderation and moderated mediation models. The index of the moderated mediation (ab) is significant for confidence intervals that do not include 0. All significant values are bolded. Predictor: baseline, moderator: between baseline and follow-up, mediator and outcome: follow-up. Pos memory = positive memory specificity, Neg events = Negative life events, Neg cognitions = Negative self-cognitions, Dep sympt = Depressive symptoms. Levels of the moderator are 0 (no events) and 1+ (one or more events). Path a1/a2 = conditional effect of predictor on mediator, b = relationship between mediator and outcome, ab = indirect effect of predictor on outcome, through mediator, c’ = direct effect of predictor on outcome controlling for the indirect effect, c1/c2 = conditional direct effect of predictor on outcome. Effect = standardised coefficient, S.E. = bootstrapped standard error, df = degrees of freedom, 95 % CI = 95 % bootstrapped confidence interval of the estimate.

| Path | Predictor | Moderator | Mediator | Outcome | Effect | S.E. | df | t | 95% CI | P(>\|z\|) |
| --- | --- | --- | --- | --- | --- | --- | --- | --- | --- | --- |
| c1 | Pos memory | 0 events |  | Neg cognitions | -1.150 | 1.232 | 418 | -0.934 | [-3.571, 1.271] | .351 |
| c2 | Pos memory | 1+ events |  | Neg cognitions | -6.530 | 1.500 | 418 | -4.353 | **[-9.479, -3.582]** | **.001** |
| a1 | Pos memory | 0 events | Neg cognitions |  | -0.773 | 1.200 | 418 | -0.644 | [-3.132, 1.585] | .520 |
| a2 | Pos memory | 1+ events | Neg cognitions |  | -5.968 | 1.463 | 418 | -4.080 | **[-8.843, -3.092]** | **.001** |
| b |  |  | Neg cognitions | Dep sympt | 0.583 | 0.044 | 419 | 13.370 | **[0.497, 0.668]** | **.001** |
| ab | Pos memory | Neg events | Neg cognitions | Dep sympt | -3.026 | 1.290 | 419 |  | **[-5.752, -0.704]** |  |
| c' | Pos memory | Neg events | Neg cognitions | Dep sympt | 0.265 | 0.858 | 419 | 0.309 | [-1.422, 1.951] | .758 |
| a1 | Pos memory | 0 events | Dep sympt |  | -0.466 | 1.286 | 418 | -0.362 | [-2.995, 2.062] | .717 |
| a2 | Pos memory | 1+ events | Dep sympt |  | -2.772 | 1.568 | 418 | -1.768 | [-5.855, 0.310] | .078 |
| b |  |  | Dep sympt | Neg cognitions | 0.513 | 0.038 | 419 | 13.370 | **[0.438, 0.589]** | **.001** |
| ab | Pos memory | Neg events | Dep sympt | Neg cognitions | -1.184 | 1.167 | 419 |  | [-3.630, 0.962] |  |
| c' | Pos memory | Neg events | Dep sympt | Neg cognitions | -2.133 | 0.799 | 419 | -2.670 | **[-3.703, -0.562]** | **.008** |

In this study, we find a dual processing effect of positive memory specificity on both cognitive and physiological vulnerability for depression in at-risk adolescents. We further reveal a potential cognitive mechanism whereby specific positive memories buffer against a tendency toward lower negative self-cognitions in response to stress. Specific positive memories may help form boundaries to the scope of negative self-cognitions, thereby preventing the emergence of depressogenic symptoms^36^. We recently showed that emphasising the value of positive social experiences as part of a brief psychological treatment programme can lead to depressive symptom reduction on par with existing treatments in depressed adolescents^37^. Encoding of current positive social experiences may increase both the availability of specific positive memories and the probability of positive memories being retrieved later in life, disconfirming negative self-cognitions arising from low mood.

We propose that positive memory specificity is an adaptive mnemonic mechanism that may be especially relevant in adolescents at risk for depression. Early adverse experiences confer risk in part because being recurrently told ‘you are worthless’ and/or ignored are associated with the emergence of negative self-cognitions^38^. These comprise a cognitive vulnerability to depression which is ‘activated’ in the face of stress^21^, leading to subsequent low mood. Early adversities have also been found to alter activation of brain areas involved in the specification of positive memories (i.e., reduced hippocampal activation), suggesting a neural substrate of lower positive memory specificity after early life stress^39^. Here, we find that positive memory specificity may act as a naturalistic defense against the negative cognitive consequences emerging from new incoming stress in at-risk adolescents.

Our findings further conceptually replicate and extend findings that positive memory recall lowers acute cortisol and mood responses to stress induction in the laboratory, where mood improvements were particularly seen in resilient individuals^4^. This conceptual replication is important given calls to triangulate research findings with multiple methods and lines of evidence^40^. Our effect of positive memory specificity on depressive symptoms was dependent on exposure to stressful events as they occurred naturally over time. This conditional effect is in line with findings in a recent longitudinal community study, which did not find an association between low memory specificity and subsequent depression; however, the study did not take the potential interaction with recent life events into account^41^. Importantly, effects of positive memory specificity on negative self-cognitions during low mood and morning cortisol were only present over time, and the mechanisms involved are unknown. Our results complement research finding a delayed symptomatic and morning cortisol reduction after positive attentional bias modification training^42^. The effect of a positive memory specificity and/or attentional bias may unfold over time by regulating responses to new life events. Here, we found this to be true for negative self-cognitions and depressive symptoms. Positive memory specificity may similarly dampen cortisol responses to everyday hassles over time. Compared to such everyday stressors, the negative life events measured here may have been too infrequent to affect the relationship between positive memory specificity and morning cortisol^43^.

We have previously demonstrated that in this sample, high morning cortisol predicts conversion to major depression only in boys with high subclinical depressive symptoms^17^, and similar results have been obtained in adolescent girls^27^. Here, we find that positive memory specificity is associated with reduced morning cortisol over time, thus potentially regulating an important physiological vulnerability marker of depression (note that this effect is present for both genders; see Supplementary Results). Together, these findings suggest that positive memory specificity in adolescents who are at risk, but not yet clinically unwell, may reduce depressive vulnerability associated with elevated morning cortisol levels. Furthermore, this physiological pathway to depressive vulnerability appeared to be relatively distinct from our measure of cognitive vulnerability, which was unrelated to cortisol in the path model (see Figure 1). This dissociation is in accordance with recent research findings, where pharmacological blockade of the Hypothalamic-Pituitary-Adrenal (HPA) axis stress response had no influence on subjective mood and self-esteem responses to stress^44^. Thus, while recent theory suggests that negative biases and cortisol may be interlinked in depression^28^, we find a dissociation of cognitive and physiological vulnerability to depression in this study. Positive memory specificity may alleviate depression through distinct pathophysiological mechanisms. As of yet unidentified, intermediate neural pathways may link these mechanisms. Reward-related neural circuitry may be a promising candidate, which is related to both mood and cortisol reactivity, and is activated during positive memory recall and facilitates resilient responses to stress^4^.

Currently, we do not know precisely how positive memory specificity operates to reduce cortisol levels over time in the developing adolescent. However, there is some evidence to support a potential mediating role of reward processing in the effects of positive memory recall on mood and cortisol^4^. Blunted reward processing arising from the striatum is one of the strongest effects of early life stress on the developing adolescent brain^45^. The intrinsically rewarding properties of positive memories (where activation of the striatum underpins rekindling of positive emotion) may be lowered in depressed individuals^46^, possibly as a consequence of blunted striatal responses to reward in major depression^47^. Thus, the protective effect of positive memory specificity in these at-risk individuals may be in part due to successful engagement of corticostriatal reward circuits. The amygdala, hippocampus and ventral striatum may be particularly important in regulating the HPA axis due to their direct connections with the paraventricular nucleus, which regulates signals to the HPA axis^48^. Lower daily cortisol output is associated with sustained corticostriatal activation to positive stimuli^49^, and decreased amygdala signal coupled with increased ventromedial prefrontal activation during emotion regulation^50^. Thus, improved reward and positive emotion processing may lead to lower morning cortisol levels. Updating of reward-based learning over time through the activation of positive memories could further explain our findings of longitudinal, but not cross-sectional, relations between positive memory specificity and morning cortisol.

In a striking homology, stimulation of positive memory engrams reduced stress-induced depression-like behaviour in preclinical mouse models^5^. Optogenetic reactivation of positive memory engrams in the dentate gyrus triggered the reward system, including parts of the striatum and the amygdala. Importantly, reactivation of engrams which encoded the memory of a positive experience (i.e., meeting a female mouse), but not simple exposure to the positive situation, lowered depression-like behaviour in male mice. This suggests that recalling specific positive memories, with concurrent activation of neural systems involved in emotion and reward processing, may facilitate resilient responses to stress^51^. This benefit of positive emotion and reward activation was additionally supported by a recent neurofeedback study where the effect of positive memory recall on depressive symptoms was mediated by increased amygdala activity after training^52^. In sum, recalling specific positive memories may rekindle positive emotion and regulate cortisol output over time. The possibility that this effect is mediated by reward processing should be investigated in future research.

Positive memory specificity may be a resilience factor that facilitates adaptive responses to stress. An international consortium recently proposed a resilience framework where resilience is defined as *‘The maintenance or quick recovery of mental health following an adverse life event or a period of adversity’*^7^. In this framework, stable pre-existing factors (resilience factors) facilitate resilient responses to future stress. These are distinguished from resilience mechanisms, which reflect adaptive responses to stress. Our findings suggest that positive memory specificity comprises a pre-existing resilience factor^16,17^ that confers adaptive responses to stress (lower negative self-cognitions after negative life events; the resilience mechanism). This process may in turn lead to the maintenance or quick recovery of mental health (i.e., lower depressive symptoms) after stressful life events.

Notably, we showed no cross-sectional relation between memory specificity and both negative self-cognitions and depressive symptoms. These findings are in accordance with the resilience framework, which suggests that resilient outcomes can only be measured after some form of life stress^7^. Depressive vulnerability was stress-emergent in this study; negative self-cognitions and, indirectly, depressive symptoms were only lowered by positive memory specificity in the presence of at least one negative life event. This is in line with an emerging animal literature finding hormonal, neural and epigenetic adaptations to experimental stress, which facilitate future beneficial outcomes^53^. Based on this literature, it has been suggested that the process underlying resilient responses to stress is dynamic and interacting rather than a stable property of an organism which can be measured in a cross-sectional manner^53^. Our findings could be explained by similar adaptive processes over time, and support a dynamic conceptualisation of resilience.

Our findings have important clinical implications, as positive memory specificity is malleable and suitable for secondary prevention in at-risk populations. That is, training in recall of specific positive memories may lower risk of developing depression. Such training has already shown promise^54^. For example, real-time amygdala neurofeedback during positive memory recall improved positive memory specificity and in turn lowered depressive symptoms after training^52^. Training may address the disturbed specificity and vividness of positive memory recall observed in depressed and recovered individuals (hampering the experience of “reliving” positive memories and thereby its mood-repairing effects)^12^. A recent study of positive memory enhancement training which emphasised specific positive memory recall provided preliminary support for this hypothesis. This study found higher memory specificity and higher perceived ability to “relive” positive memories after training, improving mood in depressed individuals^55^. The mechanistic role of negative self-cognitions in our study indicates that in particular, training in accessing specific self-affirming positive memories^56^ may prevent the development of depressive symptoms. Thus, our findings may be translated into early-phase trials seeking prevention of depression onset and relapse in at-risk populations.

The current findings should, however, be interpreted with the caveat that we did not have strict experimental control over the studied variables, thereby limiting the causal inferences that can be drawn. However, the use of longitudinal path modelling allows probabilistic conclusions about causality. That is because temporal precedence (the most important criterion for causality in the absence of experimental manipulation^57^) can be established. All our analyses took baseline measures into account, together with important confounds, which strengthens the inferences that can be made. Furthermore, we conceptually replicate findings from an experimental study^4^, which provides a foundation for our inferences about causal direction. Changes in morning cortisol attributable to positive memory specificity may be interpreted as potentially causal, because we established strong longitudinal measurement invariance of the cortisol assessments. However, we cannot fully discount the alternative causal explanation that cortisol moderated positive memory specificity^58^.

There are also some methodological limitations to consider. The relatively low number of cue words (i.e., 12) in the Autobiographical Memory Test may have reduced the reliability of the measure, particularly as responses to positive and negative cue words were analysed separately. It should further be noted that as only current and not previous psychopathology was among the exclusion criteria, it is possible that ‘scarring’ effects from previous episodes of psychopathology affected the results. However, this issue is limited by that participants were recruited in early adolescence, before the age of onset of many depressive disorders^59^. Moreover, the pattern of results did not differ in individuals who were diagnosed with major depression at follow-up (see Supplementary Results). Furthermore, exploratory analyses showed that the effects of positive memory specificity were independent of those of self-esteem and rumination (see Supplementary Results). However, it should be noted that there may be other confounding variables underlying these associations (e.g., a general positive processing bias) not measured in this study.

A limitation of the cortisol sampling protocol was that cortisol was assessed at 08.00 am with a variable time interval from waking across four mornings at baseline and follow-up. However, if the measure was highly variable due to confounding from awakening times, the latent factor of morning cortisol would be expected to reflect state characteristics and not be highly stable over time. This was not the case, as morning cortisol showed strong longitudinal measurement invariance (see Supplementary Results).

A final caveat is that in the exploratory moderated mediation models, the mediator and outcome variables were assessed at the same time. However, if shared measurement variance fully explained the mediating role of negative self-cognitions on depressive symptoms, one would assume to find a significant mediation when the variables were interchanged. Yet, depressive symptoms did not mediate the effect of positive memory on negative self-cognitions at follow-up. Similarly, participants reported both negative life events in the last 12 months and depressive symptoms in the last two weeks at the same time point at follow-up, possibly inflating their (small to moderate) interrelation. This may have been affected in part by recall bias, where participants with high depressive symptoms may have overestimated the occurrence of recent negative life events. However, negative life events were ascertained in a validated semi-structured interview with particular emphasis on reducing recall bias, showing high parent-child and panel agreement in previous reports^60^. Also, any time-invariant recall bias was taken into account by controlling for baseline reporting of negative life events. Finally, the moderated mediation analyses were exploratory, and need to be replicated in independent samples. With the above caveats in mind, we tentatively suggest that lower negative self-cognitions may comprise a cognitive mechanism through which positive memory specificity decreases vulnerability to depression in response to negative life experiences.

In sum, we show that positive memory specificity is associated with lower morning cortisol and negative self-cognitions during low mood over time in at-risk adolescents. We propose that positive memory specificity comprises a resilience factor in at-risk adolescents, by moderating cognitive and physiological pathways to depressive vulnerability after life stress. Our findings replicate and extend previous experimental work^4^, showing the potential role of positive memory specificity in regulating responses to stressors as they occur naturally over time. These findings have important clinical implications, highlighting the role of remembering specific positive life experiences in adolescent mental health resilience.

## Methods

The analyses were carried out on data from the Cambridge Hormones and Mood Project^6^. We used a subsample of participants with data available for all measures (n = 427), and these did not significantly differ from the full sample (n = 575; see Supplementary Table 1). The exclusion criteria were: current mental illness, current medical illness, pervasive developmental disorders, history of epilepsy or central neurological disease or non-English speaking. Data was collected at secondary schools in the county of Cambridgeshire in the middle 1990s (see Supplementary Methods for information about recruitment). Interviews were conducted in the school setting, which increases generalisability to a context relevant for early interventions. Parents and youths gave written informed consent to join the study. The study was approved by the Cambridge Local ethics committee and was conducted in accordance with the Helsinki guidelines.

Adolescents at risk of developing depression due to high emotional temperament or exposure to early adversity were selected and followed up over 12 months. Emotional temperament was assessed with the EAS scales (Emotionality, Activity, Sociability and Shyness)^29^ completed by parents. Emotionality is associated with development of clinical depression^61^. At-risk status was defined as having at least one early risk factor, which could be: scoring high (over the 80^th^ percentile) on the emotionality scale; current marital disharmony or past breakdown; loss of/ permanent separation from a close relative or friend; history of parental psychiatric disorder; moderately to severely undesirable events in the past twelve months. Moderate to severe negative life events in the past 12 months were assessed by semi-structured interview at baseline and follow-up^60^. A clear benefit over self-report were objective panel ratings of severity, taking factors such as social context into account (see Supplementary Methods for an overview of the types of events).

The Autobiographical Memory Test (AMT)^30^ was developed to assess the content of memories evoked by an experimental cued recall procedure. The AMT is validated and shows good psychometric properties in young adolescents^62^. Participants were presented with one of six positive and six negative cues at a time (e.g., ‘happy’) and instructed to recall a specific episode in relation to that cue. 60 seconds were allowed to produce a response. Memories were coded by research assistants trained by Professor Mark Williams, who created the Autobiographical Memory Test^30^. All ambiguous / uncertain codings were discussed at a consensus meeting of trained researchers and a coding was agreed upon. Inter-rater agreement, using the same scoring procedure, has previously been reported as excellent (99.3 % for categorical responses)^13^. Specific memories were defined as an episode with a specific time and place lasting no longer than a day. Responses were coded as categorical if they referred to repeated events. We used the ratio of specific to categorical responses to positive and negative cues in our analyses.

The Depressed States Checklist^23^ is a measure of negative self-cognitions and dysphoric experience during episodes of low mood. Participants were asked to report how they felt when their mood went down at an occasion in the last month and rate their experience on 28 adjectives (i.e., not at all; slightly; moderately; very; or extremely) of which 14 were dysphoric mood descriptors (e.g., “sad”) and 14 assessed negative self-cognitions (implying a globally negative view of the self, e.g., “useless”). The distinct and interactive nature of these two components of dysphoric experience has been supported^23^.

The Moods and Feelings Questionnaire (MFQ) is a 33-item measure of self-reported depressive symptoms for use in children and adolescents^31^. Participants rated their symptoms over the last two weeks on a three-point Likert scale (*0 = not true, 1 = sometimes, 2 = true*). The scale has good psychometric properties (α = .91, test-retest: *r* = .84)^63^.

Morning cortisol was measured at 08.00 am at four occasions within a week after the baseline measurements (see Supplementary Methods for information about assay technique). The same procedure was followed 12 months later. Participants took samples on four consecutive schooldays and recorded their time of waking. The mean time from waking to sampling was 50 minutes. Morning cortisol is relatively stable over time in this cohort (estimated to 48-60% using latent state-trait modeling^17^), and our analyses focused on the proportion that changed over time.

Adolescents’ current mental state was ascertained with the Kiddie Schedule for Affective Disorders and Schizophrenia patient version^64^ and history of psychiatric illness was assessed by semi-structured interview with both adolescents and parents. General cognitive ability (IQ) was estimated from a short version of the Wechsler Intelligence Scale for Children–II^65^ including the block design and vocabulary subtests.

Path modelling, confirmatory factor analyses (CFA) and structural equation modelling (SEM) were carried out in RStudio version 1.0.153 using the packages *psych*^66^, *ggplot2*^67^ and *lavaan*^32^. CFA is a confirmatory latent variable technique where a theorised latent construct (‘morning cortisol’) load on separate indicators (cortisol assessments across several mornings), which also have a unique variance not accounted for by the latent factor (i.e., ‘error’; see Supplementary Figure 1). Path modelling is a more flexible and powerful extension to the regression model where directional hypotheses about linear relationships between variables positive memory specificity) and variables can be tested morning cortisol and negative self-cognitions/mood). In all models, we used a robust maximum likelihood estimator (‘MLR’). This allowed computing reliable statistics despite deviations from normality^32^. Results were validated in a structural equation model (SEM; which combines CFA and path modeling in the same model) using the Full Information Maximum Likelihood method, estimating missing values by assuming that they follow patterns in the observed data (see Supplementary Table 4). This estimation method offers conservative path estimates which are penalised for the total number of estimated parameters, effectively adjusting the model estimates for multiple testing. The path model described in the main analyses had 32 free parameters, which is above the common guideline of minimum n > 10 per parameter (n = 427)^68^.

The moderation and moderated mediation analyses were conducted in PROCESS 3.0 (model 1 and 7 respectively; processmacro.org) using IBM SPSS Statistics Version 25.0. We followed the recommendations of Hayes^35^ for these analyses, given its superior power and conceptual advantages over the traditional causal steps approach^69^. PROCESS offers computation of a single index testing the significance of the moderated mediation model, removing the need for separate significance tests of each path. Removing 37 outliers with *z*- scores ± 3 did not change any of the main findings reported (see Supplementary Tables 3 and 6 for results with outliers removed). All hypothesis tests conducted were two-sided. Effect sizes reported here (Pearson’s r) represent conservative estimates, as they were calculated based on *z* and *t* scores from the baseline-adjusted longitudinal models.

We report chi-square (*X*^2^) fit statistics, the root mean squared error of approximation (RMSEA) with its 90 % confidence interval, and standardized root mean square residual (SRMR). RMSEA of less than 0.05 and an SRMR below 0.1 implies a good fit^70^. We also report the comparative fit index (CFI) and the Tucker-Lewis index (TLI), where values of CFI and TLI over 0.95 represent good fit^70^.

## Data and code availability statement

The data and code supporting the analyses presented in this paper is available at the University of Cambridge research repository [https://doi.org/10.17863/CAM.23436] and the first author’s website (www.adriandahlaskelund.com).

## Acknowledgements

This research was funded by the Aker Scholarship (A.D.A.), the Royal Society (A.L.v.H.; DH15017, RGF/EA/180029, RGF\R1\180064) and Wellcome Trust (S.S.; 209127/Z/17/Z). I.M.G. is funded by a Wellcome Trust Strategic Award and declares consulting to Lundbeck. The funders had no role in the study design; the collection, analysis and interpretation of data; writing the report; and the decision to submit the paper for publication.

Thanks to Rogier Kievit for valuable input on the statistical analyses, and to Konstantinos Ioannidis for important input on a previous version of the manuscript. Finally, we thank the participants for their contribution to our research.

## Author Contributions

A.D.A., I.M.G and A.L.v.H conceptualised the study. All authors contributed to the study design. A.D.A. did the data analysis and drafted the paper under the supervision of A.L.v.H. S.S. and I.M.G. provided critical revisions to the manuscript. All authors contributed to and approved the final manuscript.

